# Gene expression and alternative splicing contribute to adaptive divergence of ecotypes

**DOI:** 10.1101/2023.04.22.537924

**Authors:** Peter A. Innes, April M. Goebl, Chris C.R. Smith, Kaylee Rosenberger, Nolan C. Kane

## Abstract

Regulation of gene expression is a critical link between genotype and phenotype explaining substantial heritable variation within species. However, we are only beginning to understand the ways that specific gene regulatory mechanisms contribute to adaptive divergence of populations. In plants, the post-transcriptional regulatory mechanism of alternative splicing (AS) plays an important role in both development and abiotic stress response, making it a compelling potential target of natural selection. AS allows organisms to generate multiple different transcripts/proteins from a single gene and thus may provide a source of evolutionary novelty. Here we examine whether variation in alternative splicing and gene expression levels might contribute to adaptation and incipient speciation of dune-adapted prairie sunflowers in Great Sand Dunes National Park, Colorado, USA. We conducted a common garden experiment to assess transcriptomic variation among ecotypes and analyzed differential expression, differential splicing, and gene coexpression. We show that individual genes are strongly differentiated for both transcript level and alternative isoform proportions, even when grown in a common environment, and that gene coexpression networks are disrupted between ecotypes. Furthermore, we examined how genome-wide patterns of sequence divergence correspond to divergence in transcript levels and isoform proportions and find evidence for both *cis* and *trans*-regulation. Together our results emphasize that alternative splicing has been an underappreciated mechanism providing source material for natural selection at micro-evolutionary time scales.

## INTRODUCTION

Understanding the genetic basis of adaptation is a key goal of evolutionary biology. Regulation of gene expression is the fundamental link between genotype, phenotype, and the environment and is therefore a crucial component of this puzzle. Regulatory variation is known to be a major source for adaptive evolution (Jones et al., 2012; Martin & Orgogozo, 2013; Signor & Nuzhdin, 2018; Whitehead & Crawford, 2006). However, research on this topic has primarily focused on gene expression level, i.e. variation in transcript abundance. The role other gene regulatory processes is comparatively understudied (Singh & Ahi, 2022; Verta & Jacobs, 2022).

Alternative splicing of pre-mRNA (AS) is one such post-transcriptional regulatory process— conserved among eukaryotes—that produces multiple unique mRNA transcripts (i.e. isoforms) from a single gene, thus enhancing transcriptome and proteome diversity (Petrillo, 2023). This occurs via a dynamic ribonucleoprotein complex called the spliceosome. AS can generate isoforms with novel functions, modulate transcript levels/turnover (Göhring et al., 2014; Kalyna et al., 2012), or have other regulatory impacts via truncated proteins (Filichkin & Mockler, 2012; J. Liu et al., 2013). These outcomes make AS a core regulatory mechanism capable of generating diverse phenotypes (Bush et al., 2017; Wright et al., 2022).

In plants, alternative splicing contributes to multiple developmental processes, in particular seed maturation, seed dormancy/germination, seedling establishment, and transition to flowering (Posé et al., 2013; Sugliani et al., 2010; Szakonyi & Duque, 2018; Tognacca et al., 2022). AS is also known to underlie plastic responses of plants to environmental stressors including drought, heat, cold, and salt (Laloum et al., 2018; Tognacca et al., 2022). Notably, the function of AS in stress response appears to be more significant in plants than in animals (Martín et al., 2021).

Beyond developmental and environmental cues, alternative splicing (and gene expression) have a heritable component that allows for direct contribution to adaptation and divergence. Variation in gene expression level and/or alternative splicing among populations can be due to sequence variation within or nearby that gene (*cis* regulation, e.g. promoters, enhancers, suppressors, splice sites) or variation in distantly located genes whose products diffuse to influence transcription/splicing (*trans* regulation, e.g. transcription and splicing factors) (Hill et al., 2021; Wang & Burge, 2008).

Recently, evidence for the contribution of AS to adaptation and population divergence has emerged from fish (Howes et al., 2017; Jacobs & Elmer, 2021; Singh et al., 2017), insects (Y. Huang et al., 2021), and mammals (Mallarino et al., 2017), among others. In plants, studies that examine differences in AS between populations or genotypes of the same species have mainly been limited to crops, comparing different domesticated varieties or domesticated versus wild populations (Lin et al., 2020; Ner-Gaon et al., 2007; Smith et al., 2018, 2021; Thatcher et al., 2014; Vitulo et al., 2014; Zhang & Xiao, 2018) or within model organisms (Khokhar et al., 2019; Lutz et al., 2015; X. Wang et al., 2019). Therefore, although such studies have shown alternative splicing to be important, we are only beginning to understand how it could be involved in adaptation and divergence under natural selection rather than artificial selection, particularly in plants.

To what extent does genetically based variation in alternative splicing and gene expression level contribute to local adaptation and population divergence? We investigated this question using common garden transcriptomic data from a pair of prairie sunflower (*Helianthus petiolaris fallax*) ecotypes originating from Great Sand Dunes National Park (GSD) in southern Colorado, USA. These well-studied ‘dune’ and ‘non-dune’ ecotypes represent an example of local adaptation, in this case to an extreme sand dune environment (Andrew et al., 2012, 2013; Andrew & Rieseberg, 2013). GSD is home to the tallest sand dunes in North America, which are marked by shifting sands, low nutrient availability, and intense exposure (Andrew et al., 2012). *Helianthus petiolaris* is among just a few plant species that live in the dunefield, and the divergence of dune and non-dune ecotypes is estimated to have occurred in the last 10,000 years (Andrew et al., 2013).

The dune and non-dune populations are in close proximity to each other, exchanging migrants and genes, but they are differentiated genetically and phenotypically due to selection, with low (but non-zero) survival when seeds are moved between habitats (Andrew et al., 2012; Andrew & Rieseberg, 2013; Goebl et al., 2022; Ostevik et al., 2016). One of the most divergent traits between the GSD ecotypes is seed size: the dune type has seeds that are more than twice as large as the non-dune type, on average. The latter has much higher fecundity, producing many small seeds in comparison, and these differences are maintained in a common garden (Ostevik et al., 2016; Todesco et al., 2020). Larger seeds have previously been shown to have higher emergence rates both on and off the dunes (Ostevik et al., 2016). Increased seed size is thus believed to be an adaptation that aids with seedling provisioning in the depleted sand dune environment (Ostevik et al., 2016). Other traits are less well characterized, but the dune ecotype is reported to have comparatively thicker stems, reduced branching, and faster seedling growth (Andrew et al., 2013; Ostevik et al., 2016).

Gene flow between dune and non-dune populations is high enough that their genetic divergence is close to zero across much of the genome (Andrew & Rieseberg, 2013). But selection is strong enough such that there are a few important regions of elevated divergence, containing alleles strongly associated with the dune ecotype (Andrew & Rieseberg, 2013; Goebl et al., 2022; K. Huang et al., 2020). Several of these regions were recently shown to harbor large chromosomal inversions that vary in frequency across the landscape and are associated with divergent traits and environmental variables, including seed size, vegetation cover, and NO_3_ nitrogen levels (K. Huang et al., 2020; Todesco et al., 2020). Within these inversions and other loci under strong divergent selection, analysis of functionally annotated expression and splicing variation could add insight into the molecular mechanisms of adaptive divergence between GSD sunflower ecotypes.

The GSD system thus represents an excellent opportunity to connect existing knowledge of natural and evolutionary history to patterns of variation in different forms of gene regulation. We sought to: (1) characterize genome-wide differences in alternative splicing and expression (transcript levels) between ecotypes, (2) gain a more holistic view of the transcriptomic changes underlying local adaptation by comparing patterns of gene coexpression, (3) explore how regulatory divergence corresponds to genome-wide sequence divergence, and (4) determine putative functional roles of genes experiencing divergent regulation.

## MATERIALS & METHODS

### Plant material, seedling traits, RNA extractions, and sequencing

We collected *H. petiolaris* seed from three sites in the dune habitat and three sites in the non-dune habitat of Great Sand Dunes National Park, Colorado, USA in 2017 (Fig 1a). Seeds were cold stratified and germinated on filter paper prior to planting 20 seedlings from each of the six sites. Germination and planting occurred from July 3–6, 2018. For one of the non-dune sites, only nine seedlings were planted due to limited germination. Seedlings were grown in an even mix of sand and potting soil and were kept in a greenhouse setting in Boulder, Colorado. Temperature was kept between 60–80F, humidity ranged from 20–40%, and light was natural. Plants received an approximately equal amount of water each day, to saturation. We randomly selected 12 seedlings of both ecotypes for RNA extraction and sequencing, with even sampling across sites (N = 4 per site). We chose to sample seedlings because this is a consequential life stage for GSD sunflowers, especially in the dunes (Goebl et al., 2022). On July 30, we harvested the top ~100mg of tissue from selected ~3.5 week-old seedlings, which included meristem, new leaves, and upper stem. Immediately before harvest we measured height (sand to meristem) and number of leaves of all seedlings, including those that weren’t sequenced. At this time we also removed one fully expanded leaf (not included in sampling for RNA) for measurement of leaf dry mass. After harvest, we measured total dry mass of remaining above-ground tissue for seedlings not chosen for RNA-sequencing. Plants of both ecotypes were harvested at the same time and developmental stage, with most plants at the 8-leaf seedling stage (Fig. 1C). We immediately flash-froze the tissue in liquid nitrogen, stored it at −80°C, and extracted RNA the following day using a Qiagen Plant RNA Mini Kit. The meristem, new leaf, and upper stem tissue of a single plant was disrupted together to gain enough RNA for sequencing. This pooling of multiple cell types means that we can’t determine tissue-specific expression differences between ecotypes. We expect that, as with all multicellular plant RNA-seq experiments, observed expression differences are due to changes of expression within a cell type in addition to different compositions of cell types. Library prep (KAPA mRNA HyperPrep Kit) was performed for each of the 24 samples by the CU Boulder Biofrontiers Sequencing Core, followed by sequencing on an Illumina NextSeq using a 75-cycle High Output v2 reagent kit. This produced 19–23 million reads (75bp single-end) per library.

**Figure 1.**
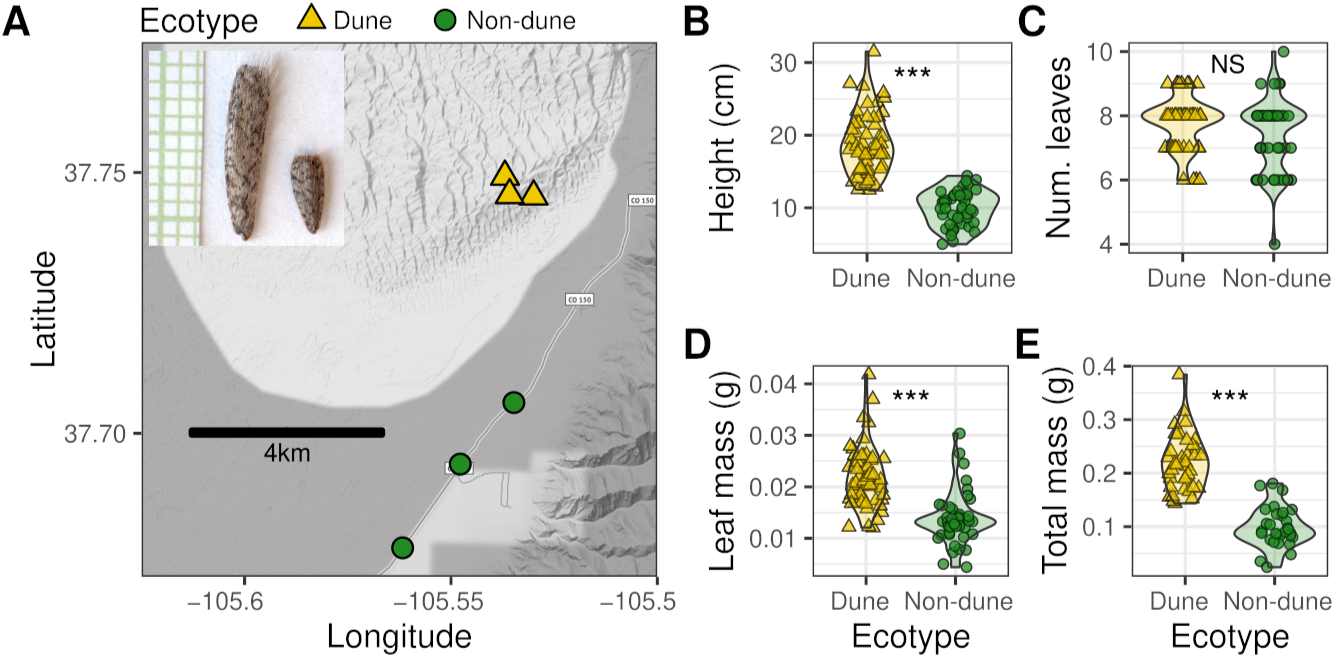
**A)** Map of sampling locations for dune (yellow triangles) and non-dune ecotypes (green circles). Inset shows typical seeds of the dune (left) and non-dune (right) ecotypes with 1mm grid paper. Photo credit Rose Andrew. **B–E)** Seedling traits measured in a common garden ~3.5 weeks post germination. Up to 60 plants of both ecotypes (20 per sampling location) were planted and measured; just 12 of both were randomly selected for the RNA sequencing experiment. Asterisks denote statistical significance of t-tests; NS, not significant. B) Height measured from sand/soil to meristem. C) Number of leaves. D) Dry mass of one fully expanded leaf. E) Total above-ground dry mass. Total mass was not measured for plants harvested for RNA sequencing.

### Filtering and read mapping

Adapter sequence and low-quality reads were trimmed using *fastp* v*0.23.1* (Chen et al., 2018). We aligned trimmed reads to the *Helianthus annuus* reference genome assembly Ha412HOv2 (Badouin et al., 2017; K. Huang et al., 2022) using STAR *v2.7.10a* in two-pass mode (Dobin et al., 2013).

### Variant calling and SNP annotation

We identified SNPs using GATK *v4.2.5.0* (McKenna et al., 2010). We first processed sorted bam files from STAR using `AddOrReplaceReadGroups`, `MarkDuplicates`, and `SplitNCigarReads`. We then ran HaplotypeCaller in `-gvcf` mode and used `CombineGVCFs` and `GenotypeGVCFs` for genotyping. We selected only bi-allelic SNPs using `SelectVariants` and applied a generic set of hard filters (`--window 35 --cluster 3 --filter “FS > 30.0” --filter “QD < 2.0”`) using VariantFiltration. We then used vcftools *v0.1.15* (Danecek et al., 2011) to filter for (1) phred quality score above 30, (2) minor allele frequency threshold of 0.05, (3) minimum read depth of at least 5 per genotype, and (4) 0% missingness. Lastly, to avoid spurious SNPs due to paralogous alignments, we counted heterozygotes per-site using vcftools *`-`*`-hardy` and filtered out sites with >60% heterozygosity.

### Analysis of sequence divergence

We calculated genome-wide *F*_st_ (Weir & Cockerham, 1984) per-site between dune and non-dune ecotypes using vcftools. We also used averaged *F*_st_ for two different window sizes: 1) 500kb non-overlapping windows for visualization, and 2) single gene windows plus/minus 5kb, which were used to investigate the association between sequence divergence and expression/splicing divergence, described below. The latter was done using the python library scikit-allel *v1.3.3* (Miles et al., 2021). Lastly, we performed a principal components analysis of filtered SNPs using the R package SNPRelate *v1.30.1* (Zheng et al., 2012). For the PCA we pruned SNPs on linkage disequilibrium with an *r^2^* threshold of 0.2, a sliding window of 500kb, and a step size of 1 SNP, using PLINK *v2* (Chang et al., 2015).

### Read counting and differential expression

Reads mapping to each gene in the reference assembly annotations were counted using HTSeq (Anders et al., 2015), which counts only uniquely mapped reads by default. We excluded genes with total read counts less than 24 as a pre-filter and then used DESeq2 to analyze differential expression (Love et al., 2014). We identified significant differentially expressed (DE) genes at FDR < 0.05 and log_2_ fold-change (LFC) > 0, following previous similar studies (Carruthers et al., 2022; Grantham & Brisson, 2018; Jacobs & Elmer, 2021; Steward et al., 2022). We also applied a shrinkage function to LFC values using `lfcShrink()` in DESeq2 in order to better visualize and rank DE genes (A. Zhu et al., 2019).To assess overall divergence in gene expression between ecotypes, we performed principal components analysis on regularized log-transformed count data (N=32,308 genes) using the R package vegan *v2.6* (Oksanen et al., 2020).

### Differential splicing

We used two approaches to analyze differential splicing between ecotypes: (1) rMATS *v4.1.2* (Shen et al., 2014), which identifies alternative splicing events using reference genome read alignments produced by STAR, and (2) the approach from Smith *et al*. (2018, 2021), which uses a custom pipeline for analyzing *de novo* transcriptome assemblies. The latter approach complements the reference-guided analysis by avoiding reference bias during transcript assembly and potentially characterizing more complex or novel splicing events.

The rMATS program is capable of detecting five major types of splice events: skipped exon (SE), intron retention (IR), mutually exclusive exons (MXE), alternative 3′ splice site (A3SS), and alternative 5′ splice site (A5SS). It counts reads that align across splice junctions and within exons to estimate the “percent spliced in” (PSI) value of each event, for each individual. PSI ranges from 0 to 1 and represents the proportion of reads mapping to one of two alternative isoforms (dubbed the “inclusion” and “skipping” isoform; importantly, while each event comprises two alternative isoforms, rMATS can identify multiple splicing events per gene, which would be expected if a gene has three or more isoforms). The degree of differential splicing for each event is then calculated as the difference in PSI between ecotypes [ΔPSI = mean(PSI*_dune_*) - mean(PSI*_non-dune_*)]. ΔPSI ranges from 1 to −1, with the extremes representing fixed differences in isoform proportions between ecotypes. By default, rMATS excludes splice events where one or both of the ecotypes have zero reads to support the event and also removes events where neither ecotype has at least one read for either the inclusion or skipping isoform. We opted to increase this threshold to require at least 12 reads in each case (up from at least 1 read) in order to increase statistical power by avoiding splice events with low support. We implemented this more conservative filter within the rMATS source code (rmats.py) because there is no option to adjust rMATS read filtering with the program’s command line arguments. We also enabled the detection of novel splice sites using the rMATS option `--novelSS`, since the Ha412HOv2 reference annotations have only a single transcript annotated per gene and divergence between *H. annuus* and *H. petiolaris* may be significant enough to alter exact splice site positions. Significant differentially spliced (DS) events were called at the default ΔPSI threshold of 0.01% and an alpha level of 0.05 after FDR correction. After significance testing we removed events that had 40% or greater missingness (N=193 events). Lastly we performed a PCA of alternative splicing using the event PSI scores, with missing PSI values imputed as the average PSI for that event.

For the Smith *et al*. differential splicing analysis, we first assembled a transcriptome for *H. petiolaris* using Trinity *v2.13.2* (Grabherr et al., 2011) with all 24 samples. We removed redundant transcripts with CD-HIT-EST at a threshold of 99% similarity (Fu et al., 2012). Next we aligned the transcriptome to the Ha412HOv2 reference genome with BLASTN and only considered hits that had at least 85% identity and aligned for at least 75% of the transcript length. We estimated isoform abundance with RSEM (Li & Dewey, 2011) and imposed the following filters for low expressed genes/isoforms: (1) we removed Trinity ‘genes’ (and thus each of its corresponding isoforms) that were only expressed in one ecotype (minimum total read count per ecotype of 24, with at least 8 samples per ecotype having at least 3 reads) and (2) we removed isoforms with total read count across all samples less than 24. Subsequent steps to filter the transcriptome, retain high confidence alternative isoforms, and test for differential splicing were the same as described previously (Smith et al., 2021). Briefly, we required alternative isoforms to align to the same genomic region, otherwise we labeled the isoforms as separate genes. We performed pairwise or multiple sequence alignments of isoforms of each Trinity ‘gene’ using EMBOSS needle *v6.6.0.0* or MUSCLE *v5.1* (Edgar, 2021), respectively, to determine if the isoforms assembled by Trinity indeed represented alternative isoforms, or if they were more likely different alleles of the same isoform. In the latter case, we clustered alleles and summed their abundance estimates (transcripts per million, TPM). Next we converted isoform TPM values to proportions of overall gene TPM values and reduced the dimensionality of each gene’s isoform composition matrix using an isometric log ratio (ILR) transformation. Finally, we tested for significant splicing differentiation between ecotypes using t-tests—or MANOVA for genes with more than two isoforms—and used an FDR threshold of 0.05 to correct for multiple testing.

To assess the congruence between rMATS results and those from the ‘Smith *et al*.’ *de novo* transcriptome pipeline, we aligned the longest isoform of each *de novo* Trinity gene, including those filtered out in the above steps, to Ha412HOv2 gene sequences using BLASTN and retained all hits with a bit score greater than 100. We generated the Ha412HOv2 gene fasta file using AGAT *v0.8.0* (Dainat, 2023) and included both intronic and exonic regions, since Trinity genes might harbor intron retention events. PCA was performed using the ILR transformations of the isoform proportions of genes with just two alternative isoforms. We did not include genes with more than two isoforms in the PCA because their isoform composition matrix remains multidimensional even after ILR transformation.

### Association between sequence divergence and expression/splicing divergence

We used a Kruskal–Wallis test to determine whether DE and DS genes differed significantly in *F*_st_ from genes that were not DE or DS; we used Wilcox tests for post-hoc pairwise comparisons. We also fit generalized linear models to assess the association between per-gene sequence divergence (see above for *F*_st_ methods) and expression/splicing divergence. For these regression analyses, expression (log_2_ fold change, LFC) and splicing divergence scores (ΔPSI) were set to zero if deemed not significant. For splicing divergence, if a gene had multiple AS events, we only used the event with largest absolute value ΔPSI. We fit zero-inflated models in both cases and specified a beta distribution to model ΔPSI ~ *F*_st_ and gamma distribution for LFC ~ *F*_st_. These models were fit with the R package glmmTMB. Pseudo-*R^2^* values were estimated using the R package performance.

### Tests of DE and DS overlap and spatial enrichment

We performed hypergeometric tests to determine the significance of overlap between DE and DS gene sets, using the R function `dhyper()`. The representation factor of the overlap was calculated as the observed divided by the expected number of overlap genes, where the expected overlap is the number of DS genes times the number of DE genes divided by the total number of genes expressed in our experiment.

We counted DE and DS genes within and outside of four major inversion regions: pet5.01, pet9.01, pet11.01, and pet17.01, using BEDTOOLS *v2.26.0* (Quinlan & Hall, 2010). We focused on these regions out of seven previously identified putative inversions (K. Huang et al., 2020) because they were by far the most divergent between ecotypes in our dataset (Fig. S1) and have been shown to contribute to adaptation in the dunes (Goebl et al., 2022; K. Huang et al., 2020; Todesco et al., 2020). While other inversions exist and are segregating, they are not strongly divergent between populations (K. Huang et al., 2020; Todesco et al., 2020). We performed Fisher’s exact tests with the R function `fisher.test()` to determine whether these inversion regions as a whole were enriched for DE or DS genes.

### Proximity of divergently regulated genes to previously identified loci under selection

A recent study of the GSD sunflowers imposed experimental selection on GSD sunflowers planted in the dunes and measured change in allele frequencies from pre- to post-selection (Goebl et al., 2022). We labeled the top 5% or the top 1% of these SNPs according to allele frequency change in hybrid (dune x non-dune) plants grown on the dunes as loci under selection/adaptive loci (see Goebl et al. 2022 Figure S10B) and subsequently tested whether DS and DE genes from our present study are more proximal to these loci compared to the null expectation, using the same approach as in (Verta & Jones, 2019). The null expectation is derived from the the proximity of adaptive loci to repeated random samples of non-divergently regulated genes. We note that tests of proximity like this focus on regulatory loci operating in *cis*.

### Investigation of putative *cis* and *trans* splicing regulatory loci

We annotated the filtered SNPs with SnpEff v5.1 (Cingolani et al., 2012), which identifies putative functional impacts e.g. splice site variants. We also identified putative spliceosomal genes in the Ha412HOv2 assembly based on homology (tblastn/blastn e-value threshold 1e-20) to *Arabidopsis thaliana* core spliceosome components and other splicing-related genes obtained from KEGG and arabidopsis.org.

### Gene coexpression network analysis

We created signed, weighted gene coexpression networks for each sunflower ecotype with the R package WGCNA *v1.71* (Langfelder & Horvath, 2008). Here we used a more stringent read count filter to reduce noise in the networks, keeping genes with mean of at least 10 reads per sample and with zero reads in no more than 6 samples of either ecotype; this resulted in 24,421 genes. We used rlog-transformed count data for input into WGCNA. The networks were built using the function `blockwiseModules()` with all replicates of a particular ecotype (N = 12 in both cases) and with a soft-thresholding power of β=18 to achieve a good model fit for scale free topology. Modules (groups of highly interconnected genes) were defined using hierarchical clustering and the dynamic tree cut algorithm with a minimum size of 30 genes. Similar modules were merged at a cut height of 0.25, corresponding to a correlation of 0.75. The remaining function parameters were set to the default values, e.g. Pearson correlation was used for construction of the networks. Separately, we used iterativeWGCNA (Greenfest-Allen et al., 2017) with default settings (except: power = 18; minModuleSize = 30) to check the robustness of coexpression networks obtained from the standard WGCNA network construction described above.

We compared coexpression networks between dune and non-dune ecotypes using the `modulePreservation()` function from WGCNA (Langfelder et al., 2011). This is a differential network analysis method that assesses the extent to which connectivity patterns among nodes (genes) of modules in a reference network are maintained in a test network. For each module it calculates mean and variance of seven network-based preservation statistics related to density (i.e. the extent to which genes retain strong connectedness) and connectivity (i.e. the similarity of intramodular connection patterns). Permutation is used to construct random modules and estimate a null distribution for each statistic, from which P-values and Z-transformed scores are calculated (Langfelder et al., 2011). The Z-scores of each metric are then aggregated into a composite summary score, the preservation “*Z_summary_*”. A *Z_summary_* score > 10 indicates strong preservation, 2 < *Z_summary_* < 10 indicates weak to moderate evidence of preservation, and *Z_summary_* < 2 indicates no evidence of module preservation. These thresholds were recommended by Langfelder *et al*. (2011) based on simulations. For this analysis we set the non-dune network as the reference network.

### Gene ontology analysis

Gene ontology annotations are lacking for the Ha412HOv2 genome, so we based our GO enrichment analyses on *Arabidopsis thaliana* GO annotations (ATH_GO_GOSLIM.txt) downloaded from arabidopsis.org (Berardini et al., 2004). We first matched Ha412HOv2 transcripts to Araport11 *A. thaliana* peptides (Cheng et al., 2017) using blastx with an e-value threshold of 1e-20, retaining only the top hit for each transcript. Next, we merged the list of *H. annuus*–*A. thaliana* homologs with the *A. thaliana* GO association file to annotate Ha412HOv2 genes with GO terms. We then used the python package GOATOOLS (Klopfenstein et al., 2018) to perform Fisher’s exact tests of GO term enrichment, for DE genes (separately for dune up-regulated and non-dune up-regulated sets) and rMATS DS genes, plus the genes that were both DE and DS (DE∩DS). We used the complete set of expressed genes (total read count >= 24, N=32,308) and expressed multi-exonic genes (N=28,389) as the background gene lists, respectively, and an FDR threshold of 0.05 for significance. We clustered redundant GO terms from each enrichment test using GOMCL (Wang et al., 2020).

## RESULTS

### Seedling traits

Despite plants being harvested at the same time and developmental stage according to leaf number (dune mode 8; non-dune mode 8; dune mean 7.9; non-dune mean 7.2; t-test p = 0.08; Fig. 1C), dune ecotype seedlings were nearly twice as tall, had over twice the mass, and also had larger leaves compared to the non-dune ecotype on average (t-tests, p<<0.001, Fig. 1B,D,E). These patterns were maintained when only considering the 24 plants randomly selected for RNA sequencing (results not shown).

### Variant calling and sequence divergence between ecotypes

The amount of uniquely mapped reads per library (i.e. per plant) ranged from 72% to 84% and averaged 80%. Un-mapped reads averaged 5.9%; the remaining reads mapped to multiple loci. Variant calling with GATK HaplotypeCaller produced 3,007,876 variable transcriptomic sites across the Ha412HOv2 reference genome, which includes 17 chromosomes plus unplaced contigs. These variants were filtered to 295,383 high quality bi-allelic SNPs. Subsequent pruning of SNPs that were in linkage disequilibrium resulted in 29,113 independent loci. PCA of LD-pruned SNPs showed distinct clustering of ecotypes along the first PC, which explained ~9.3% of the overall SNP variation among samples (Fig. 2A).

**Figure 2.**
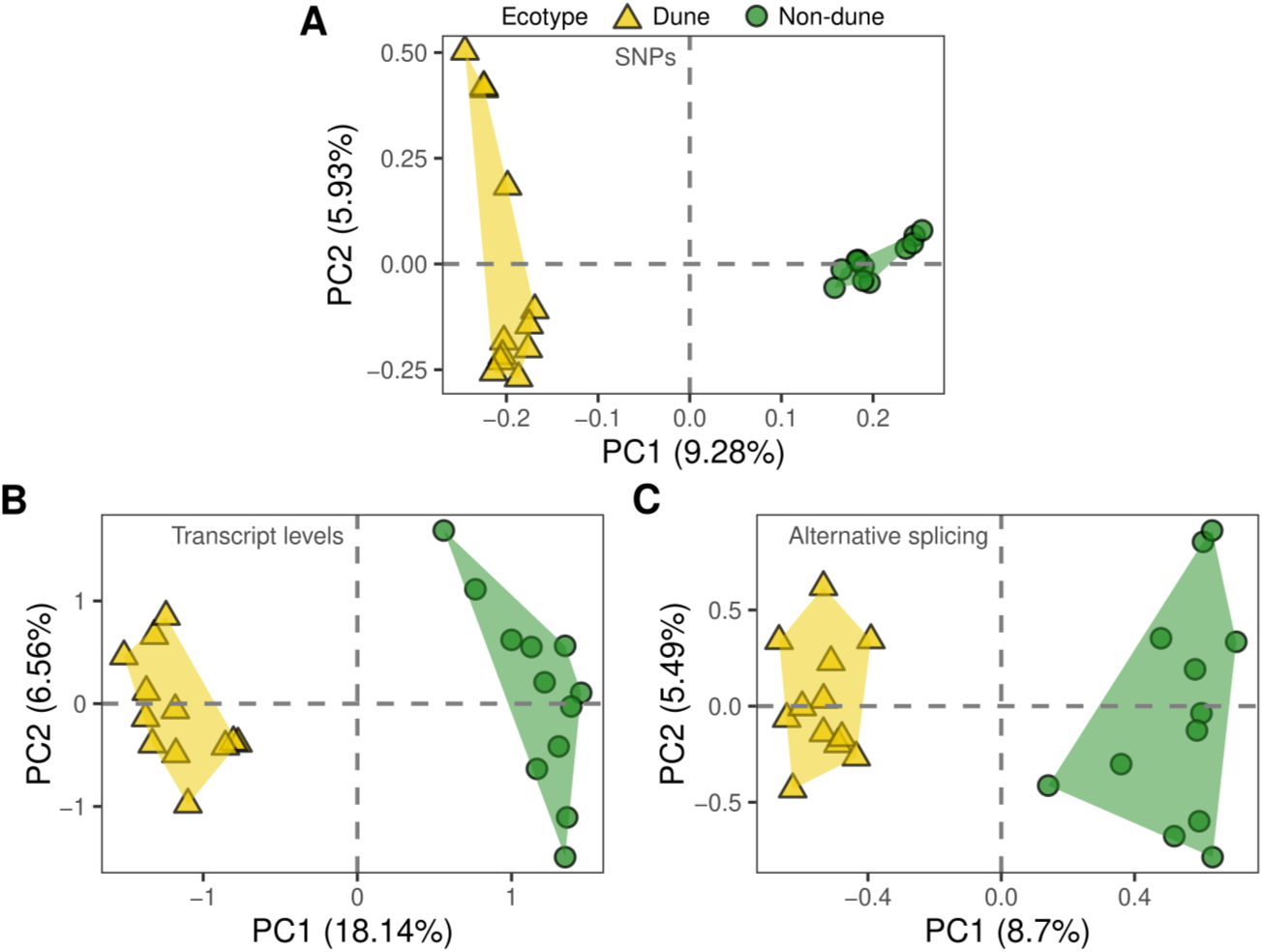
Sunflower ecotypes cluster according to sequence and regulatory variation. **A–C)** Principal component analyses showing variation along the first and second principal components for **A)** 29,113 LD-pruned transcriptomic SNPs **B)** transcript levels of 32,308 genes and **C)** isoform proportions of 17,845 alternative splicing events identified with rMATS. Percent of total variation explained by each PC is given in parentheses.

### Gene expression divergence between ecotypes

RNA-seq libraries from bulk seedling tissue produced detectable expression for 32,308 genes, representing roughly 70% of the total genes annotated in the Ha412HOv2 genome. PCA based on the transcript levels of these genes clearly separated dune and non-dune ecotypes along the first principal component, which explained 18.14% of the total variation (Fig. 2B). We found significant differential expression between dune and non-dune ecotypes for 5,103 genes (|Log2FC| > 0, FDR < 0.05), approximately 15% of genes expressed in our study (Extended Data S1). Of these, 2,480 were up-regulated in the dune environment, while 2,623 were up-regulated in the non-dune environment.

### Alternative splicing divergence between ecotypes

We detected 17,845 alternative splicing events across 6,551 genes using rMATS (Extended Data S2). This means approximately 23% of the 28,839 multi-exonic genes expressed in our study showed evidence of alternative splicing. Intron retention was the most common event type (9,946 RI events, ~55%), as expected for a plant species. PCA of isoform proportion values (percent spliced in, PSI) of all AS events produced separation between ecotypes along PC1, similar to what was found for SNPs and transcript levels, with the first axis explaining 8.7% of the total variation (Fig. 2C).

We identified 1,442 differential splicing (DS) events among 1038 unique genes (rMATS ΔPSI > 0.01%, FDR < 0.05), which represent around 16% of alternatively spliced genes. Intron retention remained the most prevalent event type among the significant DS events, though to a slightly reduced extent (669 significant RI events, ~46%). The dune ecotype tended to retain more introns (360 events with positive ΔPSI) compared to the non-dune ecotype (309 events with negative ΔPSI). Exon skipping was the least frequent AS event type overall but had a comparatively higher frequency in the set of DS events (7.9% vs 13.7%, Fig. S2).

Compared to rMATS we obtained similar results with the Smith *et al*. (2018, 2021) *de novo* isoform-based analysis of alternative splicing. We identified 6,050 Trinity ‘genes’ with clear cases of alternative splicing, and 75% of these genes had strong BLAST hits to the 6,551 rMATS AS genes (Fig. S3A). Of the 6,050 Trinity AS genes, 1,281 were significantly differentially spliced (DS), representing a similar fraction compared to rMATS (1038 DS / 6,551 AS). The 1,281 Trinity DS genes had high confidence BLAST hits (bit score > 100) to approximately 28% of rMATS DS genes (290 of 1038, Extended Data S2). Considering only reciprocal best BLAST hits, the overlap between rMATS and Trinity DS genes was 53 genes, representing the highest confidence examples of differential splicing (Extended Data S2). The small overlap in DS genes between rMATS and the Smith *et al*. pipeline is similar to what has been reported between other tools (Mehmood et al., 2020), and we think is reasonable because they differ in a number of ways: pipeline A (rMATS) tests significance of DS for a single AS event at a time, i.e. whether a particular intron or exon in spliced in or out, whereas pipeline B (Smith *et al*.) uses abundance estimates (TPM) of whole isoforms and can test for DS among three or more isoforms; pipeline B uses the ILR transform and many other steps. PCA of isoform proportions for Trinity AS genes with just two isoforms (4,050 of 6,050) again showed distinct clustering of ecotypes along the first principal component (Fig. S3B), further supporting the results from rMATS.

### Gene coexpression network divergence between ecotypes

We observed substantial differences in patterns of gene coexpression between ecotypes, which can provide insight into how the transcriptome is evolving as a whole. Coexpression networks for dune and non-dune ecotypes showed scale-free topology (Figs. S4, S5) and had similar numbers and sizes of modules (dune N = 260, non-dune N = 280; dune mean module size = 92.4 genes; non-dune mean module size = 85.3 genes). However, connectivity patterns of modules and overall network structure were not well preserved between ecotypes (Figs. 3, S5). Average *Z_summary_*preservation score of non-dune modules in the dune network was 2.2, meaning that non-dune modules had very low preservation in the dune network overall (Fig. 3). Indeed, only 7 modules showed strong preservation (*Z_summary_* > 10); 80 modules had weak to moderate preservation (2 > *Z_summary_* > 10); 195 had no preservation (*Z_summary_* < 2). Networks constructed with iterativeWGCNA had similar module numbers, sizes, and low module preservation statistics, therefore we report just the results from standard WGCNA.

**Figure 3.**
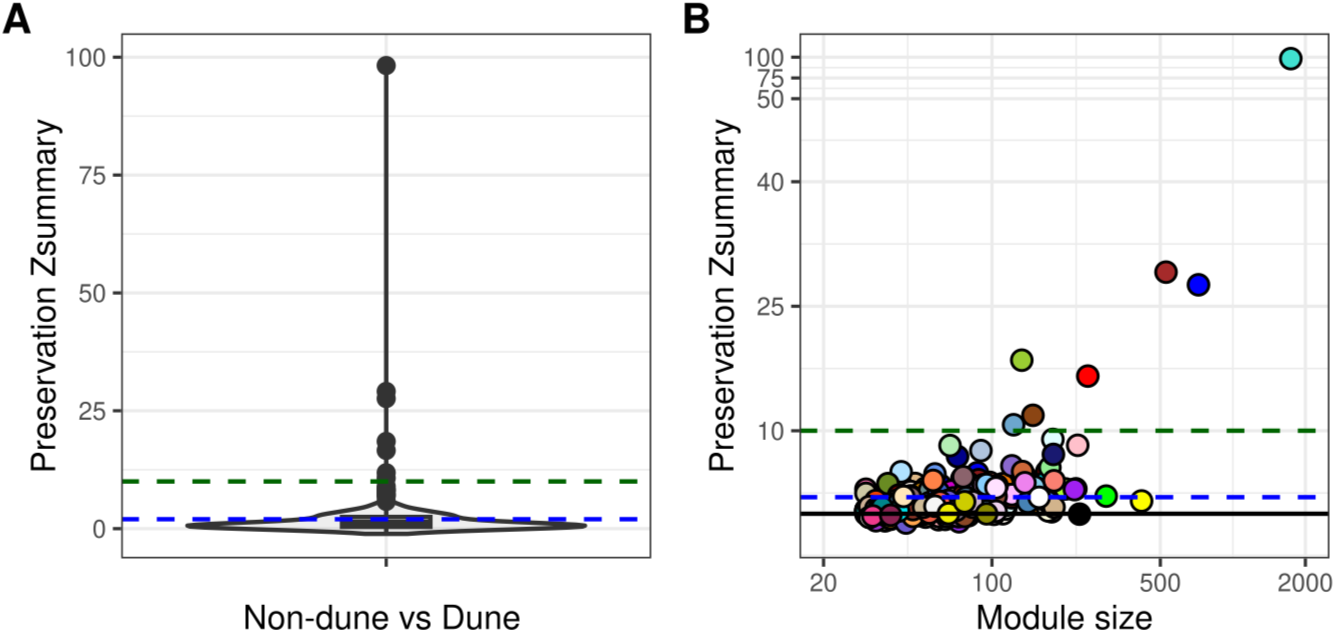
Gene coexpression networks are restructured between ecotypes. **A)** Preservation values (‘*Z_summary_*’) of non-dune network modules in the the dune network. Preservation *Z_summary_* > 10 indicates strong preservation (green dashed line); 2 < *Z_summary_* < 10 indicates weak to moderate evidence of preservation; very low preservation indicated under the blue dashed line. **B)** Module preservation of non-dune modules in the dune network, versus module size (number of genes). Colors are arbitrary module labels; similar colors do not represent inherently similar modules. The x axis is on the log scale; values 50–100 on the y-axis are squashed.

### Distribution of sequence, expression, and splicing divergence across the genome

In general, divergence in expression and splicing tended to be greater for genes in regions of higher sequence divergence, but this trend explained only a small amount of the total variation in expression and splicing differences between ecotypes.

*F*_st_ estimates in 500kb sliding windows revealed similar patterns of sequence divergence between ecotypes as reported previously (Andrew & Rieseberg, 2013; K. Huang et al., 2020; Todesco et al., 2020). We found *F*_st_ peaks in regions known to harbor large chromosomal inversions that segregate strongly between ecotypes, though some peaks existed outside these inversion regions as well (Figs. 4A, S1). This closely matches findings based on WGS data (Todesco et al 2020); previous reduced representation approaches seemingly hid some of these smaller peaks of differentiation in non-inversion regions (Andrew & Rieseberg, 2013; K. Huang et al., 2020).

**Figure 4.**
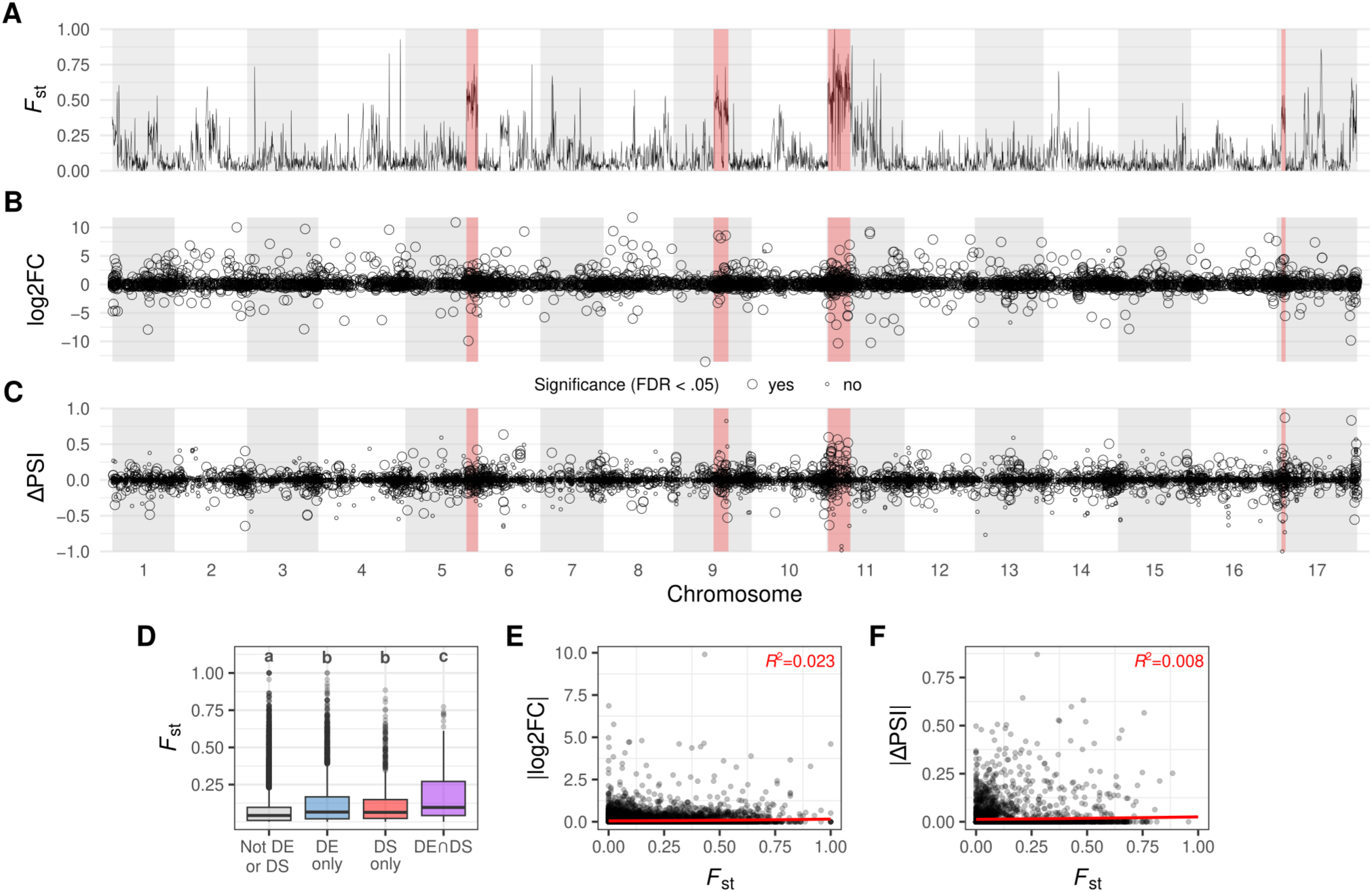
Both *cis* and *trans* regulation appear to contribute to expression and splicing divergence. Red shaded regions indicate positions of putative chromosomal inversions. **A)** Mean *F*_st_ values along 500kbp non-overlapping windows. Note putative inversion regions correspond with *F*_st_ peaks. **B)** Differentially expressed (DE) transcripts along the genome. Positive log_2_fold-change indicates up-regulation in the dune ecotype compared to the non-dune ecotype, while negative values indicate the reverse. **C)** Differential splicing (DS) along the genome. ΔPSI of 1 or −1 indicates a fixed difference in isoform proportions between dune and non-dune ecotypes. Circles represent alternative splicing events (a single gene can have multiple splice events if it has more than two isoforms). Significance of DE in **B)** and DS in **C)** is indicated by circle size, and we randomly down-sampled non-significant transcripts and events to improve visualization. Note concentration of DE transcripts and DS events within *F*_st_ peaks (e.g. pet11.01 inversion, putative *cis*-regulation) but also in regions of low *F*_st_ (e.g. chromosome 12, putative *trans-*regulation). **D)** *F*_st_ box plots for different sets of genes. Significant differences are noted with letters. **E)** Association between *F*_st_ and expression divergence per-gene. **F**) Association between *F*_st_ and splicing divergence per gene. For panels **D–F**, *F_st_* for each gene was calculated as the average of SNPs within the gene’s start/stop window ±5kb.

Divergence in transcript levels was widespread across the genome and not confined to individual regions, although some regions of high *F*_st_ also harbored numerous genes with high expression divergence, for example the inversion regions (Fig. 4B). Splicing differentiation was likewise scattered across the genome, with a notable peak of significantly differentiated splicing events within or near the pet11.01 inversion (Fig. 4C). Genes inside the four inversion loci pet5.01, pet9.01, pet11.01, pet17.01 overall were more than twice as likely to be differentially regulated (DE or DS) compared to genes outside of these inversions (Fig. S6, Fisher’s exact tests, p << 0.001). We also found that DE genes, DS genes, and DE∩DS genes tended to have higher *F*_st_ than non-divergently regulated genes (0.134 ± 0.003 DE-only; 0.124 ± 0.006 DS-only; 0.175 ± 0.013 DE∩DS; 0.083 ± 0.001 non-DE/DS; mean *F*_st_ ± standard error; Fig. 4D). Associations between *F*_st_ (single gene windows ±5kb) and log_2_ fold-change or ΔPSI were positive and significant, though weak (pseudo *R*^2^=0.023 and pseudo *R*^2^=0.008, respectively; Fig. 4E&F).

### Proximity of divergently regulated genes to loci under selection

Differentially spliced genes were consistently more proximal to previously identified SNPs experiencing the greatest allele frequency shifts following experimental selection in the dune habitat (Goebl et al., 2022) when considering distances less than or equal to 1Mb: approximately 30% of all DS genes were within 1Mb or less from a Goebl *et al*. adaptive SNP (Fig. S7A). This trend lessens beyond the 1Mb distance (results not shown), as well as when we used a more stringent threshold (99^th^ percentile) for considering SNPs as adaptive (Fig. S7B). We found that DE genes showed no significant difference from the null expectation in their proximity to adaptive SNPs (results not shown).

### Sequence divergence at splice sites and in spliceosomal genes

SnpEff annotated 884 SNPs as located in splice sites or splice regions. Average *F*_st_ of these splice variants (0.087) was comparable to that of the mean across all sites (0.084). We identified 49 splice variants among the top 5% of all SNPs based on *F*_st_, 13 of which were in differentially spliced genes (Table S1). The strongest outlier splice variant was located in the chromosome 11 inversion, had an *F*_st_ of 1, and was associated with an intron retention event in the gene Ha412HOChr11g0490131, a homolog of an *A. thaliana* organic solute transporter ostalpha protein, AT4G21570.

We found 535 sunflower genes with significant homology to known *A. thaliana* spliceosome-related genes. Average *F*_st_ of these spliceosomal homologs (0.089) was similar to that across all genes (0.095). Thirteen spliceosomal homologs were among the top 5% of all genes according to *F*_st_ and 12 of these were within one of the four major chromosomal inversions (Table S2). The splicesomal gene with strongest *F*_st_ was Ha412HOChr11g0496501, homologous to *ABH1* (AT2G13540), which encodes a nuclear cap-binding protein that is required for both pri-miRNA processing and pre-mRNA splicing and is involved in abscisic acid signaling (a key plant stress response hormone) and flowering (Cutler et al., 2010; Hugouvieux et al., 2001; Laubinger et al., 2008). The second most divergent spliceosomal gene according to sequence was Ha412HOChr11g0495951, homologous to *GFA1* (AT1G06220) which is involved in activation of the spliceosome and in embryo development (M. Liu et al., 2009; Moll et al., 2008; Zhu et al., 2016). Both these genes are within the pet11.01 inversion (Table S2).

### More overlap between DE and DS gene sets than expected by chance

More than 22% of differentially spliced genes were also differentially expressed: of the 5,103 DE and 1038 DS genes, 232 genes were both DE and DS (Fig. 5B). This overlap is significantly greater than expected due to chance, based on a hypergeometric test (p = 4.3e-09; representation factor = 1.42).

**Figure 5.**
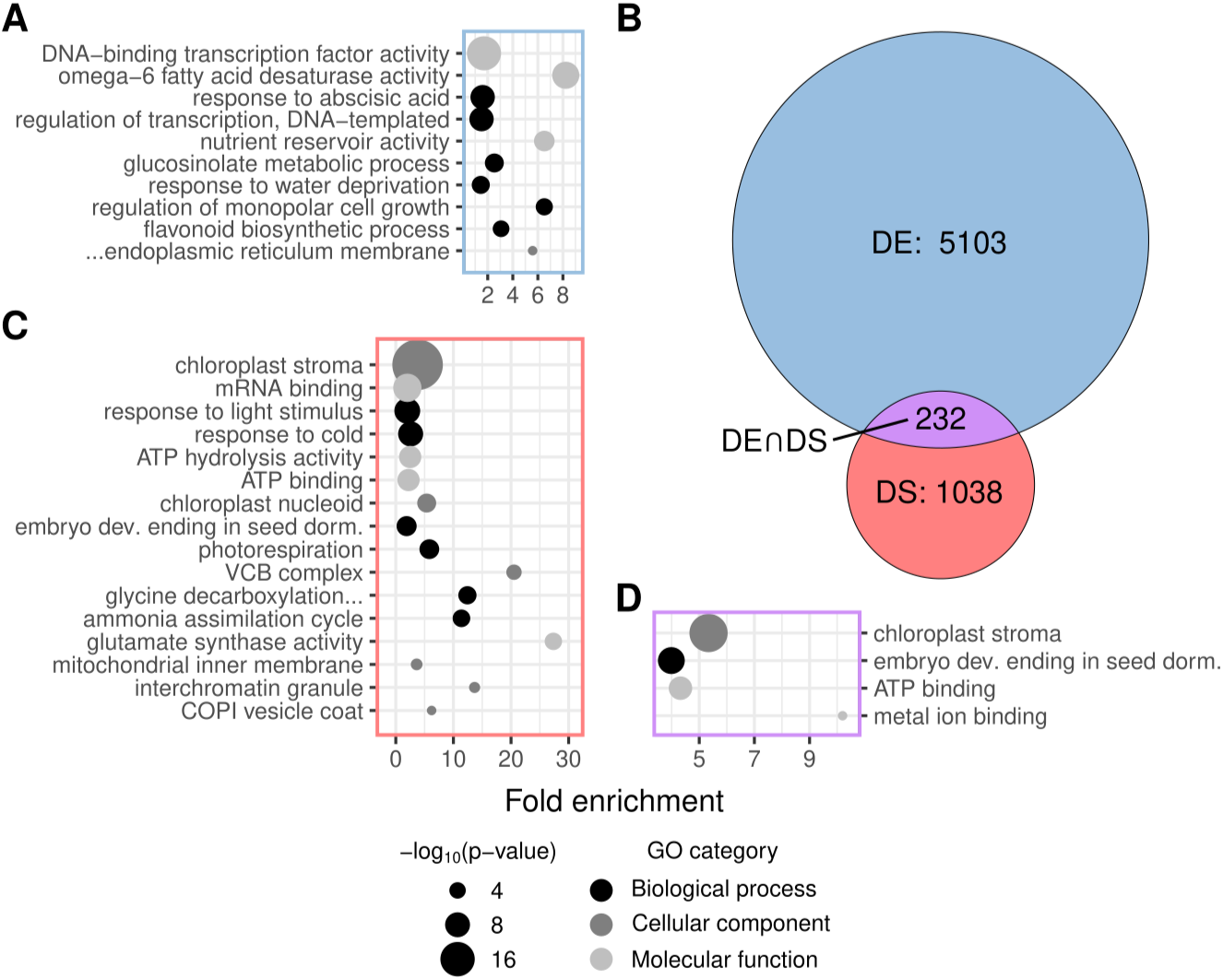
Functional enrichment of differentially regulated genes between ecotypes. **A)** GO terms enriched among DE genes up-regulated in the dune ecotype (N=2,480). **B)** Venn diagram showing number of transcripts that are differentially expressed (DE), differentially spliced (DS), or both (DE∩DS) between ecotypes (FDR < 0.05). Significance of overlap between DE genes and DS gene sets was assessed with a hypergeometric test (p < <u>0</u>.001; Representation Factor, RF = 1.4; RF values > 1 indicate more overlap than expected due to chance). **C)** GO terms enriched among DS genes (N=1038). **D)** GO terms enriched among genes that are both DE and DS (N=232). GO terms are arranged top to bottom by decreasing enrichment test significance (decreasing dot size) in panels **A**, **C**, and **D**. Some GO terms are abbreviated due to space: “…endoplasmic reticulum membrane”=“integral component of cytoplasmic side of endoplasmic reticulum membrane”; “embryo dev. ending in seed dorm.”=“embryo development ending in seed dormancy”; “glycine decarboxylation…”=“glycine decarboxylation via glycine cleavage system.”

### Putative functions of DE and DS genes

Our homology-based functional annotation (GO annotation) of the *H. annuus* Ha412HOv2 genome provided matches to *Arabidopsis thaliana* for 26,536 of the 32,3208 genes expressed in our study. We found 23 GO terms significantly enriched among genes up-regulated in the dune environment (10 after clustering), and these were primarily related to transcription, fatty acid biosynthesis, nutrient reservoir activity, and stress response processes, notably involving abscisic acid (Fig. 5A). For genes up-regulated in the non-dune ecotype, there were 152 significantly enriched GO terms (66 after clustering), and those involving translation, mRNA binding, embryo development, and photosynthetic processes/compartments were among the most significant (Fig. S8). Genes up-regulated in the dune ecotype were more likely to have a putative unique/unknown function compared to those up-regulated in the non-dune ecotype, based on fractions of either set that lacked a strong BLAST hit to *Arabidopsis* and thus lacked GO annotations (18% of dune versus 13% of non-dune up-regulated genes). This might partially explain the substantial difference in number of enriched GO terms between the two sets.

There were 34 (16 after clustering) significantly enriched GO terms among DS genes (97% of DS genes had GO annotations); these were primarily related to mRNA binding, photosynthesis, embryo development, and nitrogen assimilation (Fig. 5C). This represents an intriguing functional overlap with DE genes, specifically for those involved in embryo development and photosynthesis, which appear to be divergently regulated by both transcription and splicing mechanisms. Indeed, our GO enrichment analysis of the 232 genes that were both DE and DS recovered just four significantly enriched terms after clustering with GOMCL: chloroplast stroma, embryo development ending in seed dormancy, ATP binding, and metal ion binding (Fig. 5D).

Beyond analysis of gene ontologies, we highlight individual genes based on magnitude and significance of divergence in expression and/or splicing. Three of the top four DE genes (ranked by adjusted p-value) are nuclear-encoded homologs of ATP synthase subunit beta (*ATPB*; ATCG00480) and were up-regulated in the non-dune ecotype (Extended Data S1, Fig. S9). Homologs of four other chloroplast ATP synthase subunits (alpha, *ATPA,* ATCG00120; delta, *ATPD*, AT4G09650; gamma, *ATPC1*, AT4G04640; epsilon, *ATPE*, ATCG00470) also tended to be up-regulated in the non-dune ecotype (Fig. S9). Two homologs of *ATPI*, a subunit of the ATP synthase proton pump complex CF0, were also among the most significant DE genes, and up-regulated in the non-dune ecotype (Fig. S9).

A homolog of the *A. thaliana* splicing factor, *ATO* (AT5G06160) was the 15^th^ most differentially expressed transcript, ranked by log_2_ fold change (Ha412HOChr09g0395201; baseMean = 23, log_2_ fold change = 8.6, FDR = 1.7e-26; Fig. S10). We also identified homologs of splicing factor *SUS2* (AT1G80070) that were significantly up-regulated in the dune ecotype, with the most abundant being Ha412HOChr02g0050341 (baseMean = 213, log_2_ fold change = 1.5, FDR = 0.001; Fig. S10). Other homologs of both these splice factor genes were found in the Ha412HOv2 genome, which were not DE (Extended Data S1). Homologs of a third splicing factor, *CWC22* (AT1G80930), were also significantly up-regulated in the non-dune ecotype (Fig. S10).

Lastly we highlight one gene that overall had the third strongest splicing difference between ecotypes according to rMATS, was also DS according to the Smith *et al*. approach, and was differentially expressed: Ha412HOChr02g0088581, a homolog of *A. thaliana* glycosyl hydrolase *GLH17* (AT3G13560). This gene harbored an exon skipping event with ΔPSI of −0.644, meaning the dune ecotype more often expressed the alternative “skipping” isoform (Fig. 6). It had a reciprocal best match to a Trinity DS gene and thus is among the highest confidence examples of differential splicing (Extended Data S2). Intriguingly, the alternatively spliced (skipped) exon is in the 5’ untranslated region of *GLH17*.

**Figure 6.**
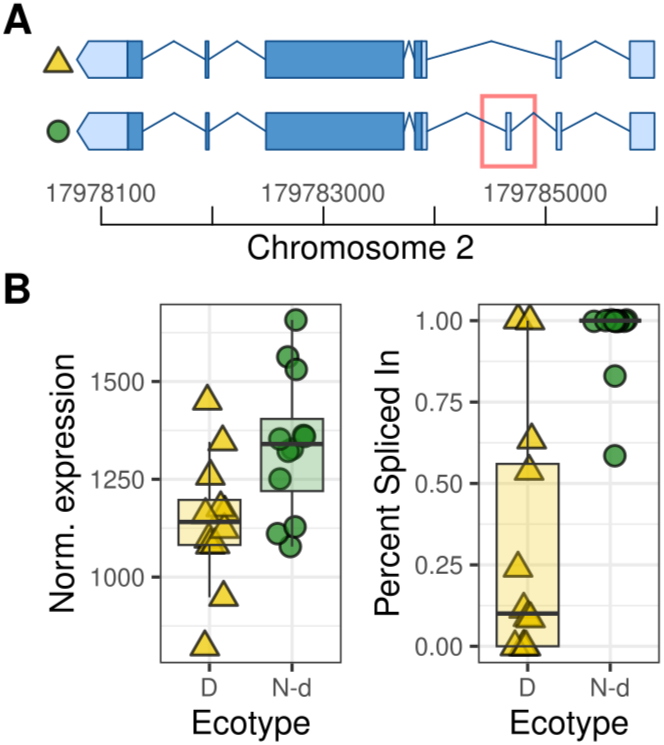
Differential regulation of *GLH17*, an example of a gene that is strongly differentially spliced and also differentially expressed between ecotypes. **A)** Structures of two alternative isoforms of *GLH17*, which constitute a case of exon skipping. The “skipping” isoform is above the “inclusion” isoform, with the skipped exon marked by a red box; isoforms are marked for whether they are more prevalent in the dune (yellow triangle) or non-dune (green circle) ecotype. Dark blue blocks represent coding exons and light blue represents exons in untranslated regions (UTRs). The alternatively spliced (skipped) exon is within the 5’ UTR. B) *GLH17 g*ene-level normalized expression differences between dune (D) and non-dune (N-d) ecotypes (log_2_ fold change = −4.3e-05, FDR < 0.05) and differences in alternative splicing (ΔPSI = −0.644, FDR << 0.001) for the exon skipping event. “Percent Spliced In” (PSI) refers to the proportion of reads supporting the inclusion isoform. Points represent individual plants.

## DISCUSSION

### Extent of expression and splicing divergence between ecotypes

Previous research has demonstrated that strong allelic differences observed between the dune and non-dune prairie sunflower ecotypes of GSD are adaptive, in other words, selection is driving allelic differences (Goebl et al., 2022). Our results show the ecotypes also have evolved distinct patterns of both gene expression levels and alternative splicing (Fig. 2B&C). Recent findings have been equivocal as to which of these two processes evolves faster at time scales involving ecotypic adaptation and to what extent they follow complementary versus independent trajectories (Carruthers et al., 2022; Jacobs & Elmer, 2021; Verta & Jacobs, 2022). The first principal component for expression level, which also delineates ecotypes, explains a substantially larger amount of overall variation compared to that of alternative splicing (Fig. 2B&C). We also observed substantially more significantly differentially expressed genes compared to differentially spliced genes (Fig. 5B), though we are cautious in taking this difference at face value because (1) the nature of short read sequencing data likely makes detection of alternative splicing more difficult than expression (2) the relative proportion of alternatively spliced genes that are DS is similar to that of expressed genes that are DE (around 15%), and (3) DE and DS are called using slightly different thresholds, in this case LFC > 0 and ΔPSI > 0.0001. Still, a similar magnitude of difference in number of DE versus DS genes exists between sexes of multiple bird species (Rogers et al., 2021), and analyses of environmentally determined phenotypes also report fewer DS compared to DE genes (Grantham & Brisson, 2018; Healy & Schulte, 2019; Steward et al., 2022). In contrast, comparisons of arctic charr ecotypes did not reveal consistently more DE than DS genes (Jacobs & Elmer, 2021).

The fact remains that expression (transcription) and alternative splicing are inherently linked. Substantial overlap in DE and DS genes, their functions, or their associated SNPs (i.e. expression and splicing QTL) has been observed in cases of plasticity (Grantham & Brisson, 2018; Healy & Schulte, 2019), ecotype differences (Carruthers et al., 2022), and species differences (Singh et al., 2017). Other studies suggest the two regulatory processes mostly evolve independently (Jacobs & Elmer, 2021; Jakšić & Schlötterer, 2016; Martín et al., 2021; Verta & Jacobs, 2022). We have shown that both transcript level and alternative splicing are associated with divergence in the GSD sunflower ecotypes. Although the majority of differentially spliced genes were not differentially expressed in our study, the ~22% overlap was larger than expected by chance (Fig. 5B). Thus there appears to be an important role of the two processes acting both independently and jointly in divergent adaptation of GSD sunflowers.

### Disruption of gene coexpression networks

Gene coexpression also appears to have been dramatically restructured in the process of adaptive divergence: strong correlations in transcript level among genes are rarely preserved when comparing the coexpression network modules of the non-dune to the dune ecotype (Figs. 3, S5). Only seven modules showed significant preservation between networks, and approximately 75% of the modules had no evidence for preservation, based on composite module connectivity and density preservation scores (Fig. 3). This result suggest that, beyond the differential expression or splicing of particular genes, novel connections among genes could be important for—or a product of—adaptive divergence. The dramatic restructuring of gene coexpression is striking, since the ecotypes are so recently diverged. Comparison of wild versus domesticated cotton also revealed substantial restructuring of coexpression networks, marked by fewer modules with tighter connections (i.e. larger modules) in the domesticated variety (Gallagher et al., 2020). A similar scenario of coexpression rewiring has been reported for maize vs teosinte (Swanson-Wagner et al., 2012). It is interesting that each of these examples, including ours, all involve recent adaptive evolutionary shifts in phenotypes, ecology, and allelic variation, and also show substantial changes in gene coexpression.

### Distribution of regulatory divergence across the genome

We were curious how gene expression level and splicing divergence is patterned across the genome in relation to sequence divergence, and what this might tell us about the relative contributions of *cis* versus *trans* regulation. We found that genes within four large chromosomal inversion regions—which correspond with high *F*_st_ peaks and have been previously implicated in the adaptive divergence of the GSD sunflowers—were overall twice as likely to be DE and/or DS compared to genes outside these regions. This indicates there are substantial *cis*-regulatory variants within these haploblocks that are contributing to splicing and expression divergence. Furthermore, DE, DS, and DE∩DS genes tended to have higher *F*_st_ than non-DE/non-DS genes (Fig. 4D), which is consistent with the influence of *cis-*regulation. Although we found only a few splice site variants with high *F*_st_ within DS genes (i.e. putative *cis*-sQTL, Table S1), we note that our characterization of specific *cis*-sQTL is very much incomplete due to technical limitations: the tool we used to annotate splice site variants does not recognize novel splice sites (of which there were many) and also does not recognize other splicing *cis* regulatory elements like splicing enhancer and silencers, which can be found further from the splice site region (Lovci et al., 2013; Wang & Burge, 2008). Lastly, we found some evidence that DS (but not DE) genes tend to be more proximal to previously identified loci under selection (Fig. S7). Because these previously identified loci were SNPs derived from reduced representation sequencing (Goebl et al., 2022), we expect that most direct targets of selection were not sequenced, and that identified adaptive loci were more likely to be indirectly affected by selection. Though we don’t have the exact targets of selection identified, the combination of these two studies suggests that divergent selection is affecting *cis* regulation of splicing such that the ecotypes express different compositions of isoforms.

On the other hand, we found widespread expression and splicing divergence outside of inversions and within low *F*_st_ regions (Fig. 4), which suggests *trans* regulatory elements in the inverted/highly divergent regions (e.g. Table S2) may modulate expression or splicing throughout the genome. We found that genes comprising transcription and splicing machinery were often differentially expressed or differentially spliced (Figs. 5A&C, S8), which is in agreement with previous findings (Jacobs & Elmer, 2021; Jakšić & Schlötterer, 2016). GO terms such as DNA transcription factor activity, regulation of transcription, and mRNA binding were enriched in either DE or DS gene sets (Figs. 5A&C, S8). A homolog of splicing factor *ATO* was one of the most divergently expressed genes between ecotypes (Fig. S10). *ATO* has been previously shown to regulate gametic cell fate in plants alongside spliceosomal component *GFA1* (Moll et al., 2008), which we found to have high sequence divergence between ecotypes (Table S2). We also found homologs of splicing factors SUS2 and *CWC22* that were up-regulated and down-regulated in the dune ecotype, respectively (Fig. S10). Alongside the *F*_st_ outlier spliceosomal homologs we identified (which included homologs of *GFA1* and *ABH1*, Table S2), these genes represent strong candidates for *trans-*regulatory loci: although their own expression and/or splicing may be controlled by *cis* variants, they could act to influence the transcription and/or splicing of genes elsewhere in the genome. Although we cannot rule out the possibility that our 500kb sliding window *F*_st_ averages are masking high divergence at individual *cis*-regulatory SNPs (Fig. 4A), regression analyses show that sequence divergence in *cis* explains only a small fraction of the total variation in expression level or splicing divergence among all genes (Fig. 4E&F), again consistent with the influence of *trans* regulation.

Together these results show that in addition to *cis* regulation, there are *trans*-regulatory variants contributing to variation in both transcript abundance and alternative splicing. We stress that future investigations would benefit from controlled crosses to map *cis-* and *trans-*sQTL (and eQTL) with more specificity, as in Smith *et. al* (2018). Still, our findings appear consistent with previous work showing trans regulatory loci tend to contribute more to expression variation within species, while *cis* regulatory divergence becomes more impactful between species (Bao et al., 2019; Schaefke et al., 2013; Signor & Nuzhdin, 2018; Wittkopp et al., 2008). We speculate that increased divergence at *trans*-regulatory loci would be expected to result in more dramatic restructuring of coexpression networks, as described above (Figs. 3, S5), since these genes often influence expression of multiple other genes [i.e. they are more pleiotropic (Vande Zande et al., 2022)].

### Divergent regulatory evolution highlights potential mechanisms of adaptation

Several traits and environmental variables differ between the GSD ecotypes/habitats over short geographic distance, including seed size, seedling size, nitrogen availability, exposure i.e. to light/temperature/wind, and soil stability (Andrew et al., 2012; Ostevik et al., 2016; Fig. 1). Of these, larger seed size and tolerance of low nitrogen levels have been most clearly implicated as adaptive traits in the dune ecotype (K. Huang et al., 2020; Ostevik et al., 2016; Todesco et al., 2020). Results from gene ontology enrichment analyses suggest gene expression and alternative splicing evolution may influence these traits. They also point towards additional genes/pathways that may contribute to adaptation in the dunes via divergent regulation.

We found embryo and seed development-related GO terms (e.g. ‘embryo development ending in seed dormancy’) enriched among both DE and DS genes (Figs. 5, S8). The expression of seed development-related genes in seedlings could be due to multiple factors, including pleiotropy and the correlation of expression across multiple tissues and developmental stages. Whether—or precisely how—divergent regulation of this category of genes gives rise to larger seeds in the dune ecotype remains unclear. Intriguingly, however, this same functional category was prevalent among genes differentially spliced between wild and domesticated common sunflower (*Helianthus annuus*) seedlings (Smith et al., 2018) and was echoed in *F*_st_ outlier spliceosomal genes, e.g. *GFA1* (Table S2). These results are consistent with developmental genes being enriched for alternative splicing (Bush et al., 2017) and implicate such genes as recurring targets of regulatory changes associated with rapid divergence.

Multiple nitrogen assimilation-related GO terms were enriched among differentially spliced genes (‘ammonia assimilation cycle’ and ‘glutamate synthase activity’; Fig. 5C), which underscores tolerance to low nitrogen availability as an important trait in the GSD system. It would be interesting to further explore how alternative splicing contributes to the underlying physiological mechanism of nutrient usage in the dunes. Adaptation to the dunes also appears to involve constitutive up-regulation of abiotic stress response genes (Fig. 5A), including those involved in ‘response to abscisic acid’, which is a central plant stress response signaling hormone (Cutler et al., 2010). Overall, the dune habitat has been shown to favor a different growth and reproductive strategy than the non-dune habitat. Consistent differences in gene expression and splicing of chloroplast and mitochondria-targeted genes between ecotypes is an intriguing pattern in this context (Figs. 5, S8, S9) and raises the possibility that cyto-nuclear interactions are an important factor in the divergence of GSD ecotypes. Photosynthesis and metabolism-related genes have been previously unexplored in the GSD system, though a recent study reported a similar pattern of down-regulation for such genes in domesticated sunflower in response to drought treatments (Lee et al., 2022). We hypothesize that divergent regulation of photosynthetic machinery and/or abiotic stress response pathways might be an additional way in which growth and stress tolerance is changed in the dunes.

As an example of a gene with strongly differentiated splicing and significantly different expression level between ecotypes, the sunflower homolog of *Arabidopsis thaliana GLH17* illustrates a way that alternative splicing may interact with expression level to affect proteome composition (Fig. 6). Specifically, our results show that this gene undergoes skipping of an exon within its 5’ untranslated region (UTR), predominantly in the dune ecotype. UTRs are well known to influence several aspects of gene expression, including rate of translation, post translational modification, and targeting of the protein (Srivastava et al., 2018); alternative splicing of 5’ UTRs likely influences translational efficiency (Chaudhary et al., 2019). *GLH17* is involved in the auxin mediated initiation of lateral root emergence (Swarup et al., 2008). Given the very different soil conditions (e.g. stability, nutrient availability) between the dune and non-dune habitats, root structure would be an intriguing phenotype to investigate further.

## Conclusion

Understanding the molecular processes that contribute to adaptation and divergence is a key goal of evolutionary biology, but alternative splicing has been understudied in this regard, with just a handful of well documented examples to date (Singh et al., 2017; Verta & Jacobs, 2022). Because alternative splicing is a important mechanism for plant development and plastic stress response, among other processes, we hypothesized it may underlie adaptive changes as well, especially in extreme habitats such as the Great Sand Dunes. We found that differences in splicing and expression level between a sand dune-adapted prairie sunflower ecotype and its neighboring non-dune ecotype were widespread throughout the genome at the seedling stage in a common environment. Overall, our results represent one of the first clear examples of genome-wide alternative splicing divergence within a non-crop plant species.

Although we do not specifically test the adaptive impact of alternative splicing and expression variation in the dunes, given that gene flow is ongoing between the ecotypes (Andrew et al., 2012, 2013), neutral divergence is expected to be limited in this system, with divergence of genes and traits driven mainly by selection (Goebl et al., 2022; Nosil et al., 2009; Yeaman & Whitlock, 2011). This raises the possibility that gene expression and alternative splicing are either indirectly or directly under selection, which is bolstered by the enrichment among DE and DS genes of functional annotations (e.g. seed development and nitrogen assimilation) seemingly related to known adaptive traits in the dune ecotype. Thus, we conclude that variation in alternative splicing and gene expression level are both likely contributing to adaptive divergence of ecotypes.

## ACKNOWLEDGMENTS

The authors would like to thank: Christopher Pauli for providing early guidance on transcriptomic analysis, Luke Evans for helpful discussion regarding data analysis, and Scott Taylor and members of the Taylor Lab for feedback on figures and an earlier version of the manuscript. Seeds were collected at Great Sand Dunes National Park under permit #GRSA-2015-SCI-0008. The CU BioFrontiers Interdisciplinary Quantitative Biology PhD program has been an important source of fellowship funding and support for PAI, AMG, CCRS, and KR (NSF IGERT grant number 1144807 and NSF NRT grant number 2022138). The authors would also like to acknowledge that GSDNP that is within the ancestral lands of the Ute and Pueblo people as well as the Jicarilla Apache Tribe and Navajo Nation.

## AUTHOR CONTRIBUTION STATEMENT

NCK conceived of the study and supervised the work. AG collected seeds. PAI and AG raised plants and performed RNA extractions. PAI performed or guided all analyses; KR performed the WGCNA analysis and drafted corresponding sections of the manuscript. CCRS provided code and assistance for the ‘Smith *et al*.’ differential splicing analysis. NCK provided guidance on all analyses. PAI wrote the initial draft and designed all figures. All authors provided feedback on the draft and contributed to revisions.

## DATA ARCHIVING

The raw RNA sequence data will be available in the NCBI Sequence Read Archive upon publication, under BioProject PRJNA996226. All code is available at https://github.com/peterinnes/Innes_et_al_2023_GSD_RNA-Seq.

