## Supplemental Figures for "Gene expression and alternative splicing contribute to adaptive divergence of ecotypes"

**Figure S1. Manhattan plot showing $F_{st}$ across the genome in 500kb windows.** Same as Figure 3A except all seven previously identified putative inversion regions are marked in red, and asterisks denote the four highlighted in the main text.

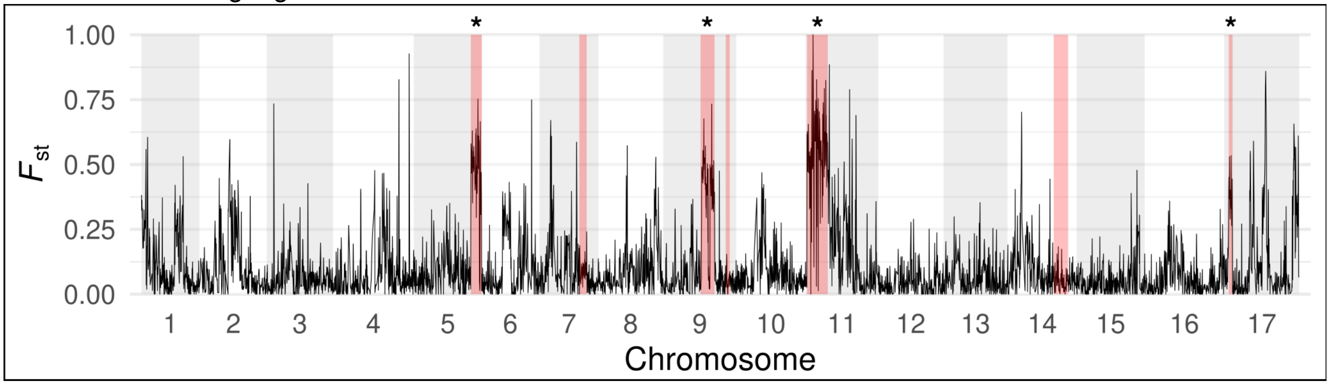


**Figure S2. Counts of alternative splicing events identified with rMATS.** RI = retained intron; A3SS = alternative 3' splice site; A5SS = alternative 5' prime splice site; MXE = mutually exclusive exons; SE = exon skipping. Significantly differentially spliced (DS) events are represented by the portion of each bar that is shaded black.

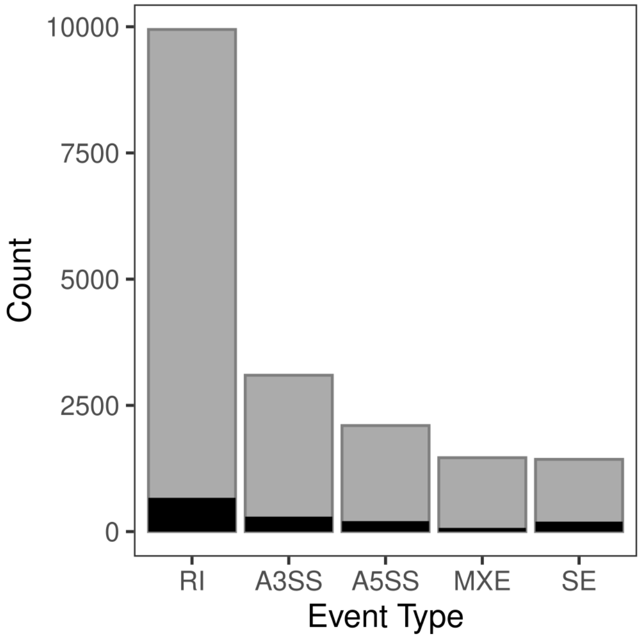

**Figure S3. Alternative splicing analysis based on *de-novo* transcriptome assembly supports results from reference-based rMATS. A)** Overlap of sets of alternatively spliced genes identified by different approaches. Approximately 75% of *de novo* assembled (Trinity) genes identified as alternatively spliced using Smith et al approach matched alternatively spliced genes found using rMATS and the Ha412HOv2 reference genome (BLASTN bit score > 100). **B)** PCA of isometric log ratio-transformed isoform proportions for Trinity genes with two alternative isoforms, as identified by the Smith et al approach (N = 4,050; the 2,000 AS genes with more than two isoforms were excluded, see Methods). Yellow triangles represent the dune ecotype, green circles are the non-dune ecotype. Compare to Figure 2C.

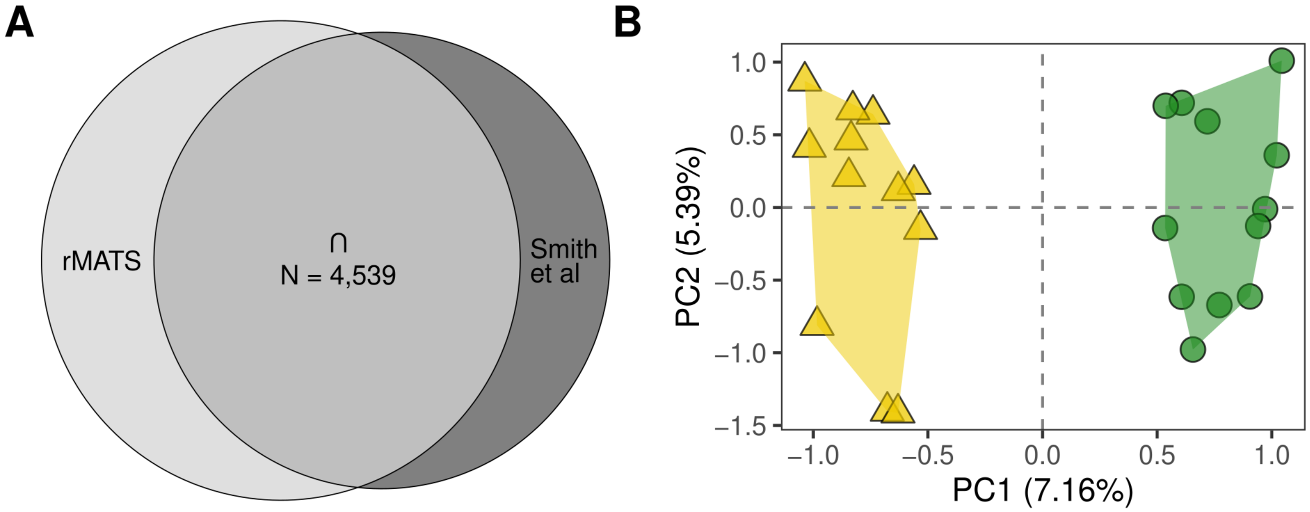

**Figure S4. Scale-free topology plots of Dune and Non-dune co-expression networks.**

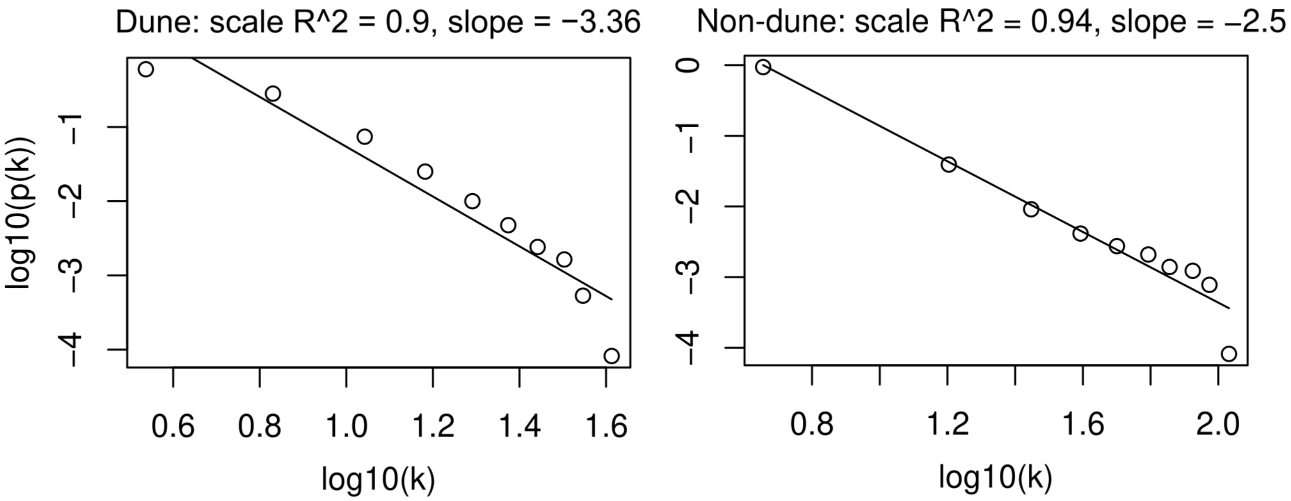

**Figure S5.** Hierarchical clustering dendrograms representing gene co-expression networks for the non-dune and dune ecotypes, built with 24,421 genes. Each branch of the tree is a gene. Non-dune module labels are shown in arbitrary color blocks. The lack of preservation of non-dune modules in the dune network is illustrated by the fact that very few blocks of color are retained when mapping the non-dune module labels onto the dune network.

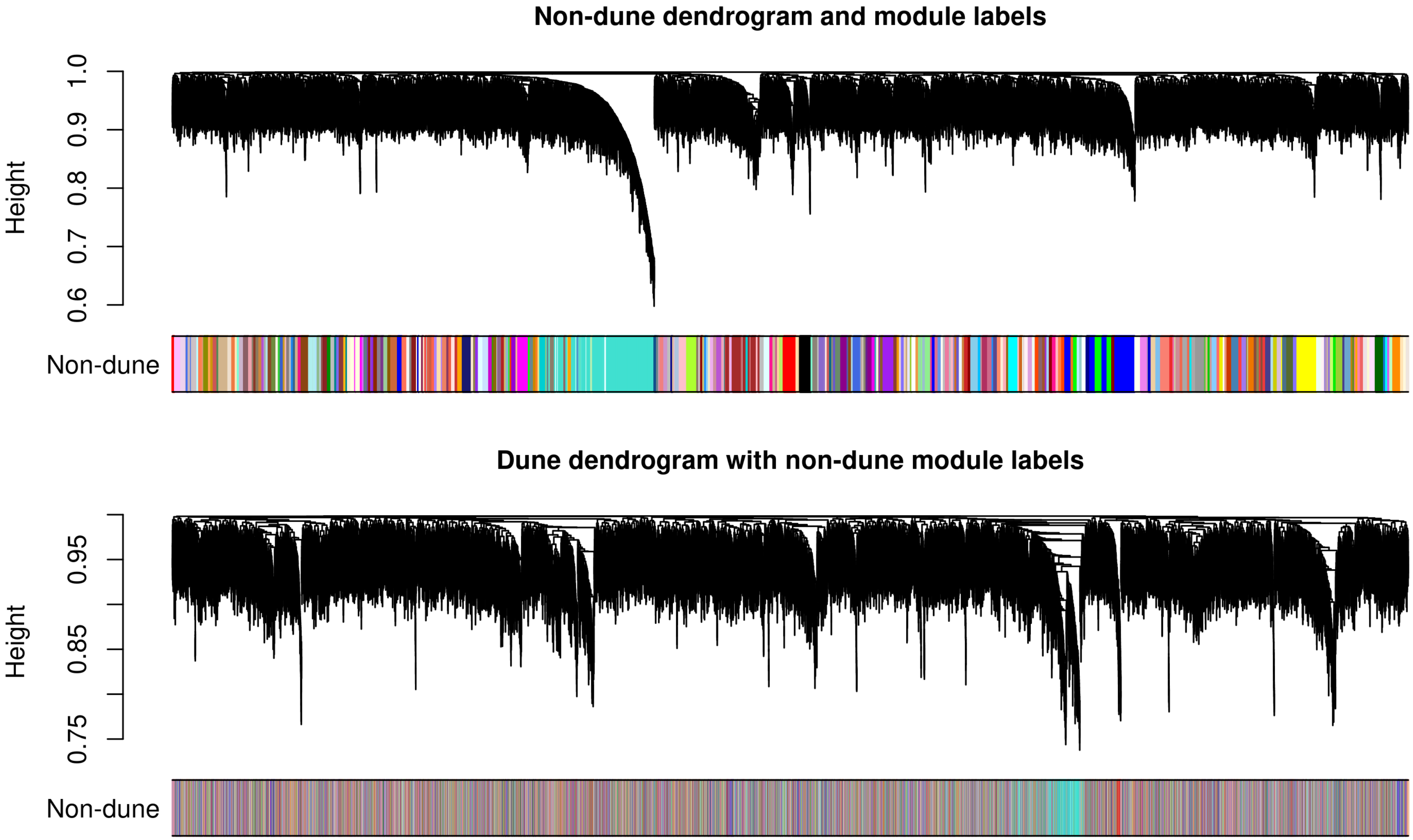

**Figure S6. Inversion regions are enriched for A) differentially expressed genes and B) differentially spliced genes.** Proportions/percentages represent the fraction of DE or DS genes within versus outside of the four inversions pet05.01, pet09.01, pet11.01, and pet17.01 (counts were combined across inversions). Fishers exact tests were used to assess the significance of enrichment (p << .001 for both tests).

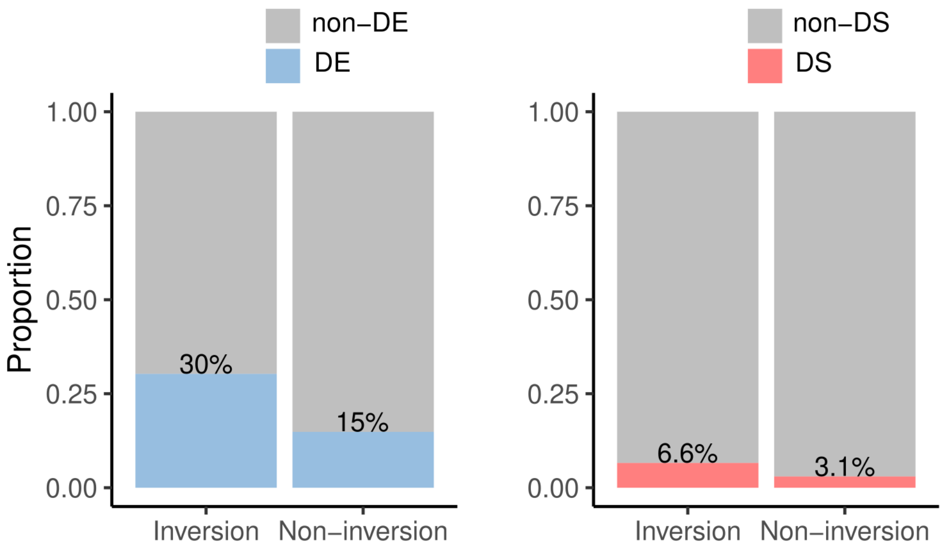

**Figure S7. Distance of differentially spliced genes to previously identified loci under selection in the dune habitat.** The red line traces the cumulative proportion of DS genes. “Control” genes (dashed line) are those not experiencing divergent regulation. Shaded region represents 95% confidence interval. Panels **A** and **B** differ in the threshold applied to designate SNPs as “adaptive” according to magnitude of allele frequency shift following experimental selection on plants in the dunes (Goebl et al., 2022). Loci (SNPs) were derived from reduced representation sequencing. **A)** Using a 95% quantile threshold, there were 611 adaptive SNPs; within a 1Mb window, DS genes we consistently closer to these adaptive SNPs compared to the null expectation; around 30% of all DS genes were located 1Mb or closer to an adaptive SNP. **B)** There were 123 adaptive SNPs using the more stringent 99% quantile threshold; in this case the signal of increased proximity of DS genes to adaptive loci is lessened.

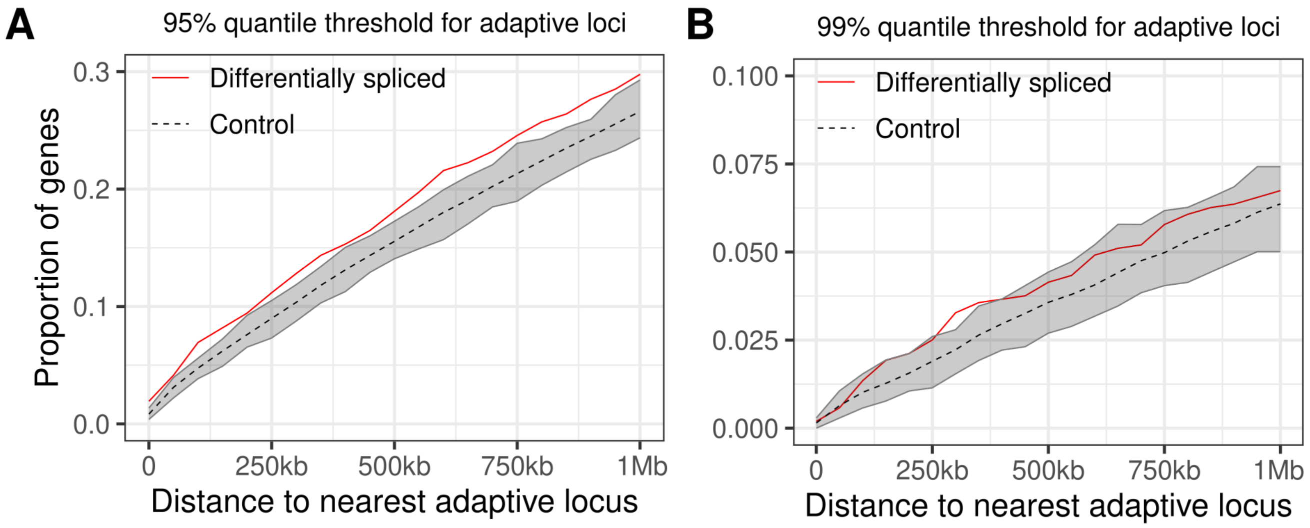

**Figure S8. Gene ontology terms enriched among 2,623 DE genes upregulated in the non-dune ecotype**, following clustering by GOMCL.

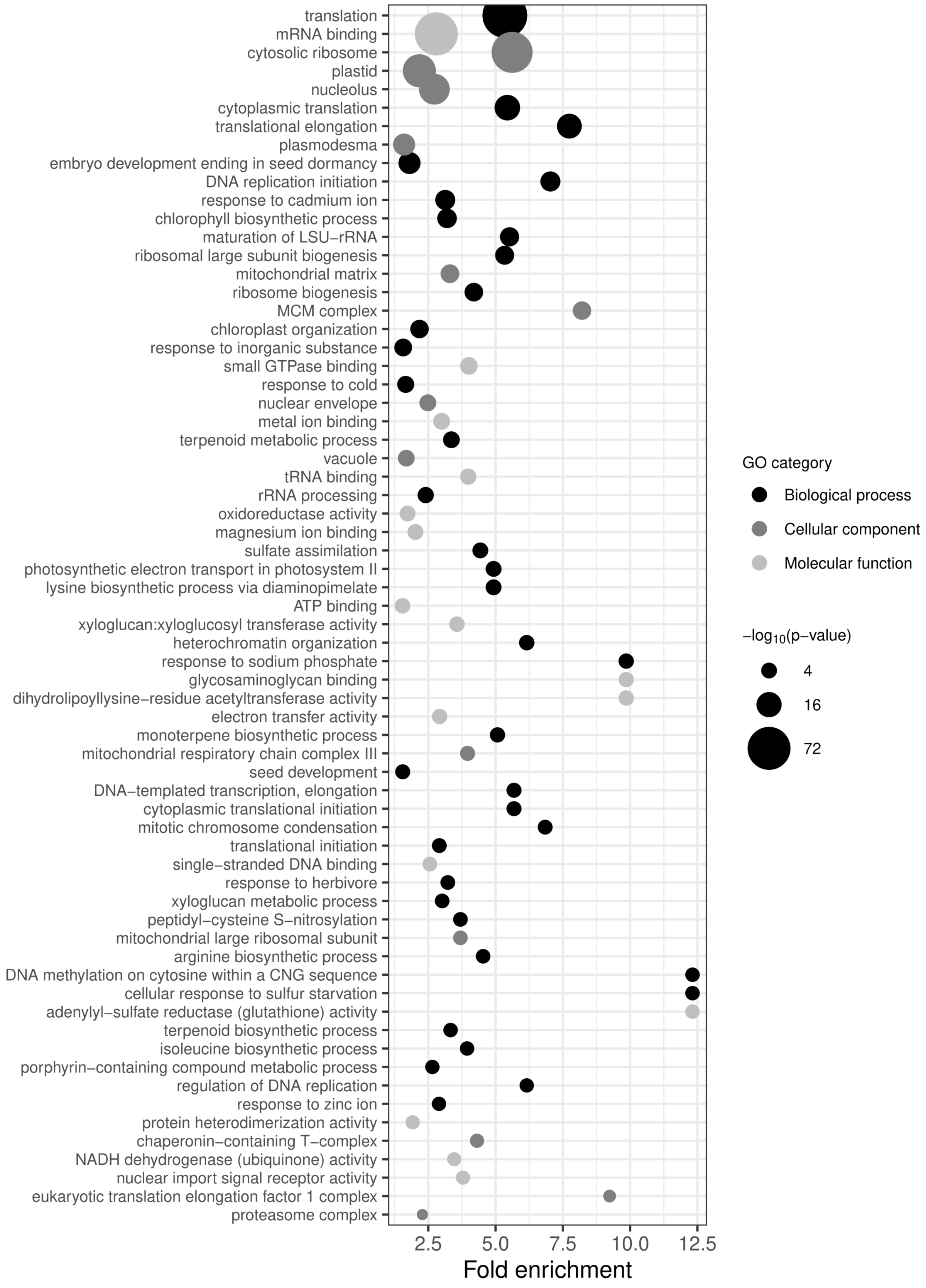

**Figure S9. Nuclear-encoded homologs of chloroplast ATP synthase subunits are consistently down-regulated in the dune ecotype compared to the non-dune ecotype.** All panels show significant differential expression (FDR < .05), except for those three labeled 'NS' (not significant). **A)** subunit alpha, ATCG00120 **B)** subunit beta,  ATCG00480 **C)** subunit gamma, AT4G04640 **D)** subunit delta, AT4G09650 **E)** subunit epsilon, ATCG00470 **F)** subunit a of ATPase complex CF0, the ATPase proton pump, ATCG00150.

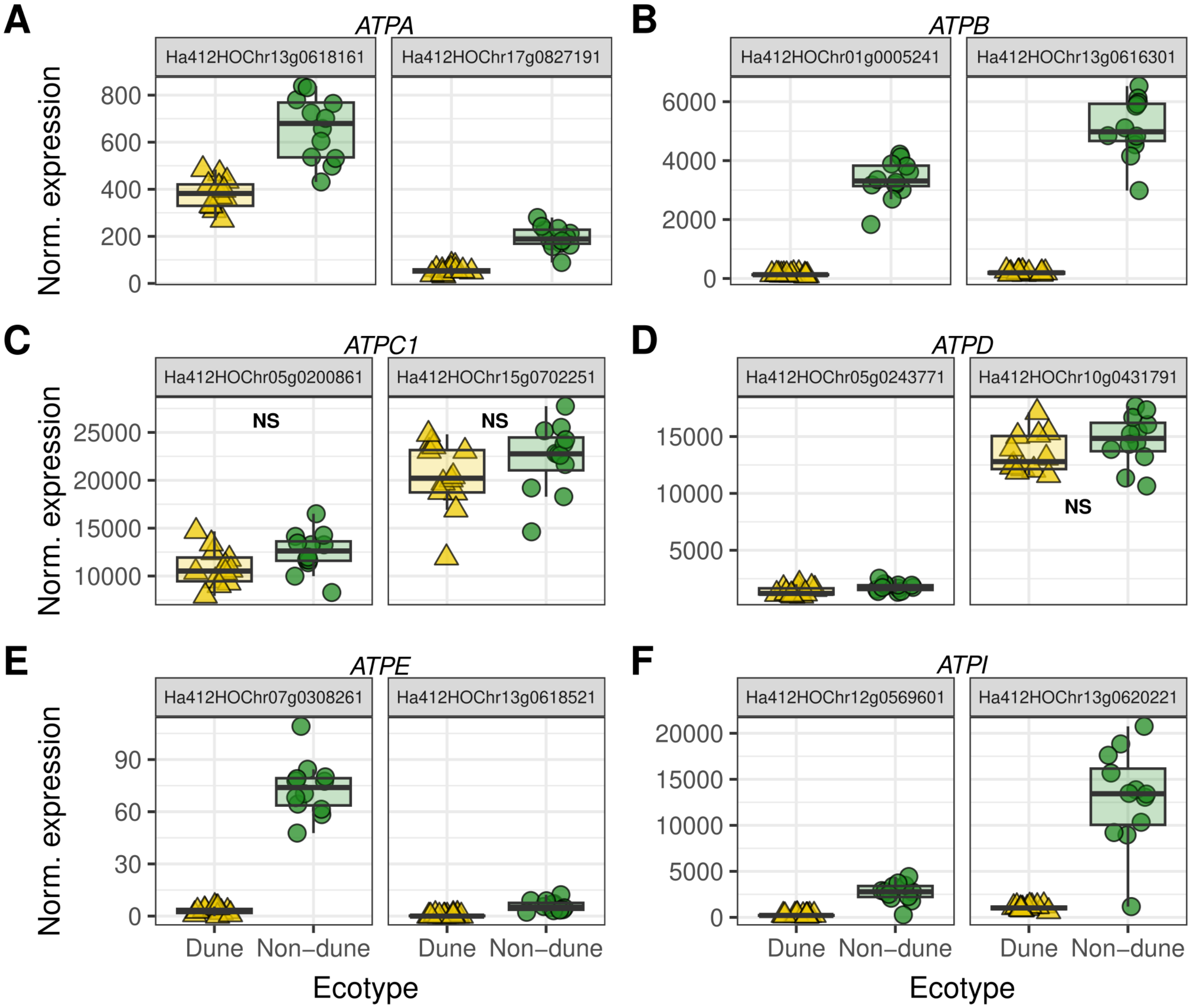

**Figure S10. Multiple splicing factors are differentially expressed between ecotypes.** Normalized expression of homologs of splicing factors **A)** *ATROPOS, ATO* (AT5G06160) **B)** *ABNORMAL SUSPENSOR 2, SUS2* (AT1G80070) and **C)** *CWC22* (AT1G80930)

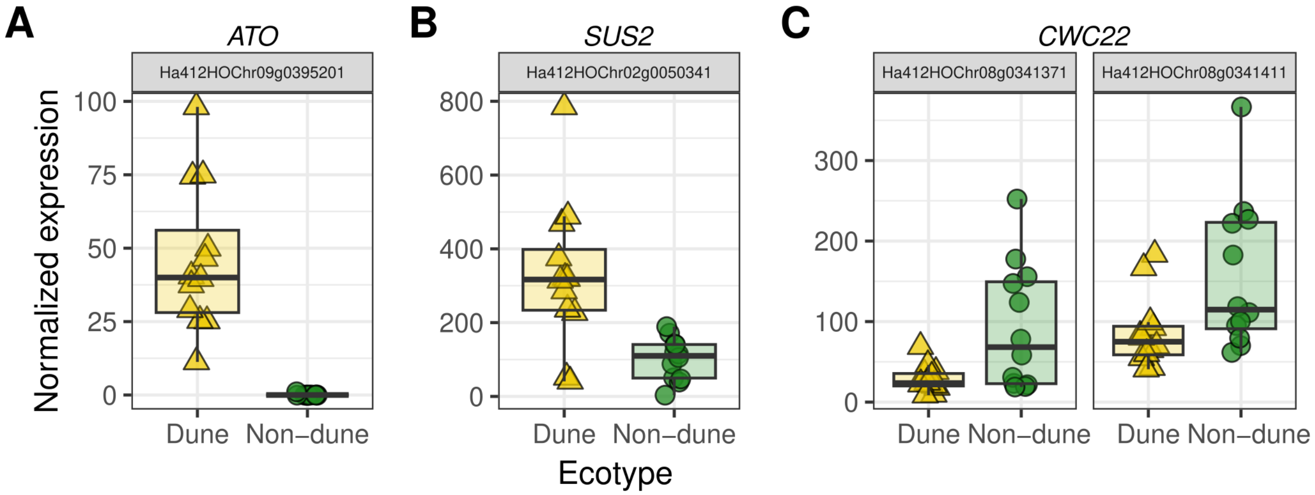
